# Modelling tuberculosis drug resistance amplification rates in high-burden settings

**DOI:** 10.1101/2021.01.12.426461

**Authors:** Malancha Karmakar, Romain Ragonnet, David B. Ascher, James M. Trauer, Justin T. Denholm

## Abstract

**Background:** Antimicrobial resistance develops following the accrual of mutations in the bacterial genome, and may variably impact organism fitness and hence, transmission risk. Classical representation of tuberculosis (TB) dynamics using a single or two strain (DS/MDR-TB) model typically does not capture elements of this important aspect of TB epidemiology. To understand and estimate the likelihood of resistance spreading in high drug-resistant TB incidence settings, we used molecular understanding to develop a compartmental epidemiological model of *Mycobacterium tuberculosis* (*Mtb*) transmission.

**Methods:** A four-strain (drug-susceptible (DS), isoniazid mono-resistant (INH-R), rifampicin mono-resistant (RIF-R) and multidrug-resistant (MDR)) compartmental deterministic *Mtb* transmission model was developed to explore the progression from DS-to MDR-TB. The model incorporated strain-specific fitness costs and was calibrated using data from national tuberculosis prevalence surveys and drug resistance surveys from Philippines and Viet Nam. Using an adaptive Metropolis algorithm, we estimated drug resistance amplification and transmission rates.

**Results:** The posterior estimates for the proportion of isoniazid mono-resistant amplification among treatment failure was 0.75 (0.64 – 0.85) for Philippines and 0.55 (0.39 – 0.63) for Viet Nam. The proportion of rifampicin mono-resistant amplification among treatment failure was 0.05 (0.04 – 0.06) for Philippines and 0.011 (0.010 – 0.012) for Viet Nam. In Philippines, the estimated proportion of primary resistance resulting from transmission was 56% (42 – 68) for INH-R, 48% (34 – 62) for RIF-R and 42% (34 – 50) for MDR-TB. For Viet Nam, the estimated proportion of drug resistance due to transmission was 79% (70 – 86) for INH-R, 68% (58 – 75) for RIF-R and 50% (45 – 53) for MDR-TB.

**Discussion:** RIF-R strains were more likely to be transmitted than acquired through amplification, while both mechanisms of acquisition were important contributors in the case of INH-R. These findings highlight the complexity of drug resistance dynamics in high-incidence settings, and emphasize the importance of prioritizing testing algorithms which also allow for early detection of INH-R.

## Introduction

Despite being both a preventable and curable disease, more than 10 million people develop tuberculosis (TB) each year, with 1.4 million deaths in 2019 [1]. Although 63 million lives have been saved through improvements in programmatic TB management this century, the increase in drug-resistant (DR-TB) cases is increasingly concerning. Multidrug-resistant TB (MDR-TB; defined as resistance to both first-line drugs isoniazid and rifampicin) is a particular barrier to TB control efforts [2]. In 2019, 465,000 people were diagnosed with MDR-TB [1]. MDR-TB can be acquired by transmission (primary resistance) or develop *in vivo* through inadequate or incomplete treatment (secondary resistance), and the relative contribution of these mechanisms is likely to vary by context [3]. In all settings, though, careful optimisation of both clinical and public health management of MDR-TB is required to ensure good outcomes.

Mathematical modelling is increasingly used to support programmatic optimization for TB [4–6]. Accounting appropriately for MDR-TB in mathematical models of disease is critical; as it differs considerably from drug-sensitive TB (DS-TB) in both epidemiological parameters and relevant outcomes. Some variation in parameters is relatively well-understood, including the prolonged treatment duration [7], adverse event rates [8] and diagnostic pathway performance [9, 10]. However, considerable uncertainties persist with regards important characteristics of MDR-TB, particularly those relating to fitness, transmissibility and risk of resistance amplification related to treatment [11, 12]. Attempts to better characterize these features of MDR-TB have been challenging, in part due to the diversity of gene mutations which may confer resistance, many of which have limited clinical and epidemiological outcome data to inform model parameterization. Computational biological approaches have recently been used to bridge this gap, providing tools to estimate the fitness and resistance impact of even novel TB mutations [13–15].

Modelling approaches also offer an opportunity to quantify amplification and transmission of drug-resistant TB, through incorporation of observed data from real-world settings. We therefore aimed to incorporate molecular data into an empirically calibrated model, in order to explore parameter estimation for drug resistance amplification and transmission associated with both isoniazid and rifampicin.

## Methods

### 2.1 Constructing the mathematical model and defining epidemiological parameters

We developed a four-strain TB transmission model to study and estimate rates of drug resistance amplification. The first strain is the drug-susceptible (DS-TB) strain (compartment subscript S), the second strain is the isoniazid mono-resistant (INH-R) strain (compartment subscript H), the third strain is the rifampicin mono-resistant (RIF-R) strain (compartment subscript R) and the fourth strain is the MDR-TB strain (compartment subscript M). We used five compartments to represent mutually exclusive health states with regards to TB infection and disease - susceptible (S), early latent (L_A_), late latent (L_B_), infectious (I) and recovered (R) to capture *Mtb* transmission dynamics. It is to be noted that the strains are not phylogenetically related.

We assumed homogenous mixing in a population of constant size N:

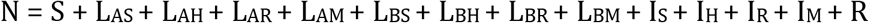

All deaths are replaced as new births (π), entering the susceptible compartment. This includes both deaths due to TB disease (μ_i_), as well as a universal population-wide death rate (μ).

When individuals in a population are infected with a circulating strain of *Mtb*, they transition from the susceptible compartment (S) to the early latent compartment (L_A_). The force of infection (λ) associated with each strain is defined as:

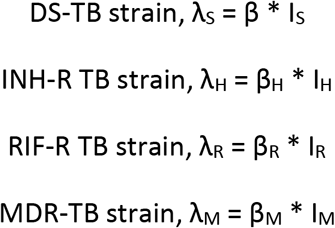

Where,

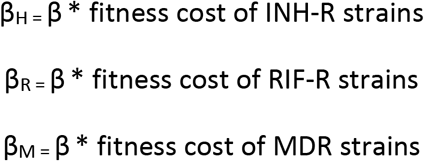

The parameter β is the “effective contact rate” for wild-type DS-TB, defined as the product of the average number of contacts between two individuals per unit time and the probability of transmission per contact.

People entering the early latent compartment (L_A_) can either progress directly to the active disease compartment (I) at rate ε, or transition to the late latent compartment (L_B_) at rate κ. Progression from L_B_ to the active disease state occurs at a much slower rate (ν), and is referred to as reactivation. Once individuals have entered the infectious compartment, one of the following can occur: 1) the person may be correctly identified as having active TB and commenced on treatment (rate τ), thence progressing towards cure and transitioning to the recovered (R) compartment; 2) spontaneously recovery (rate γ) with transition to the recovered compartment (R); 3) death (μ_i_) or 4) the infecting strain could acquire resistance (α_H_ and/or α_R_) to isoniazid (INH-R), rifampicin (RIF-R) or MDR-TB and move to I_H_, I_R_ and ultimately to I_M_ compartments. To capture progressive accrual of resistance, with each transition, only one level of additional resistance not already present can be obtained. People who have spontaneously recovered from past TB or successfully completed treatment are both represented as a single compartment (R) on the assumption that prognosis is equivalent regardless of the infecting strain from which each person recovers. Once treatment is complete, the recovered person can transition back to L_A_ through reinfection, represented as δ. We define δ as:

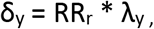

where, RR_r_ is the “relative risk of re-infection once recovered” and “y” indexes the drug resistance pattern – S, H, R or M.

Latently infected people also have a risk of re-infection with the same or other strains represented as θ in the model. We define θ as:

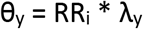

where, RR_i_ is the “relative risk of re-infection once latently infected” and “y” indexes the resistance pattern – S, H, R or M.

**Figure 1:**
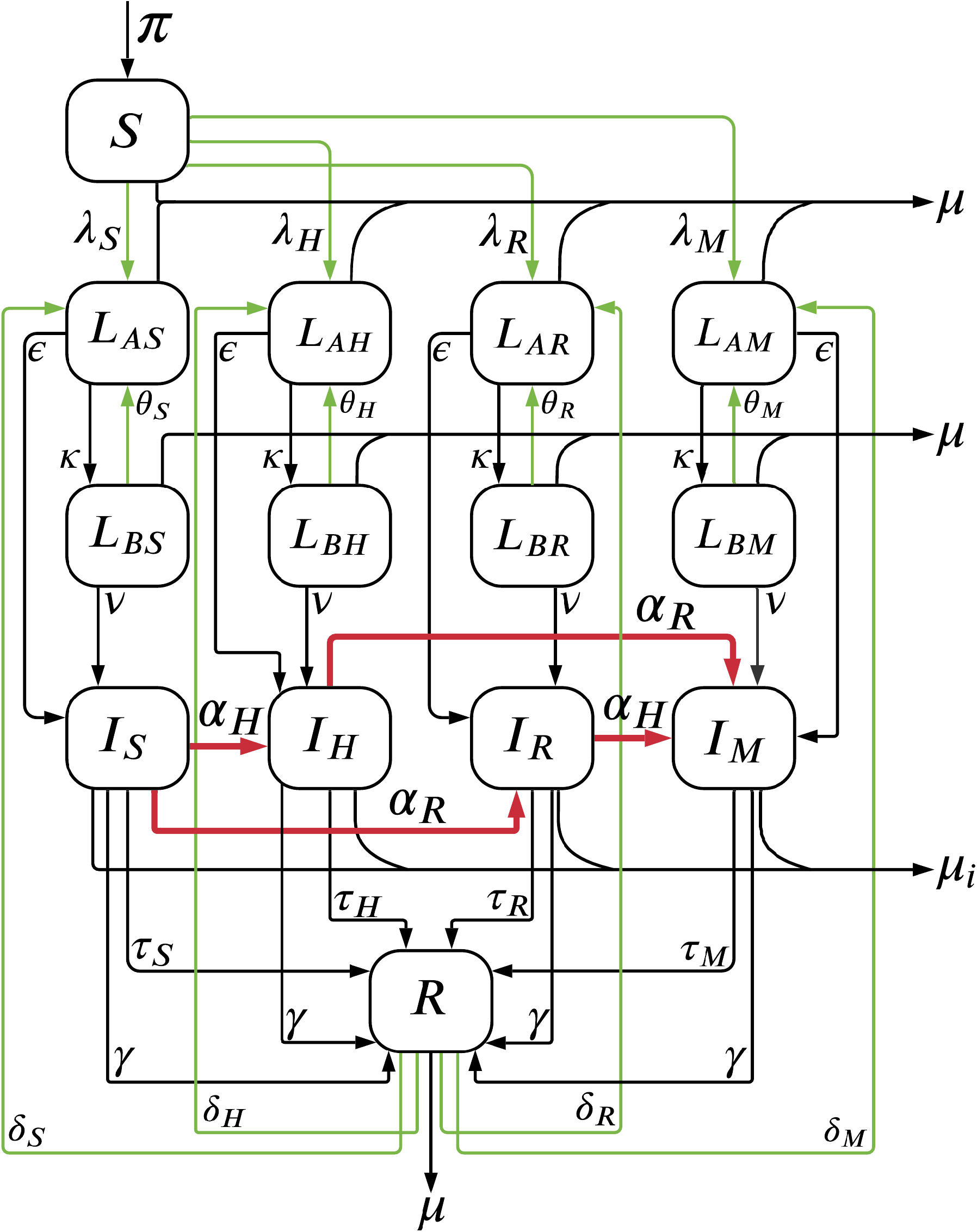
Structure of four strain TB transmission model. The symbols S, L, I and R represent uninfected/susceptible, latent, infected and recovered health states, respectively. To distinguish between various strains, the subscripts S, H, R and M are used to represent susceptible, isoniazid mono-resistance, rifampicin mono-resistance and multidrug resistance respectively. The green arrows represent infection/transmission flows, black arrows represent constant progression flows and red arrows represent amplification. (To avoid over-crowding the diagram we do not represent force of re-infection once infected (θ) with the other strains.)

#### Ordinary differential equations used to define the four-strain model

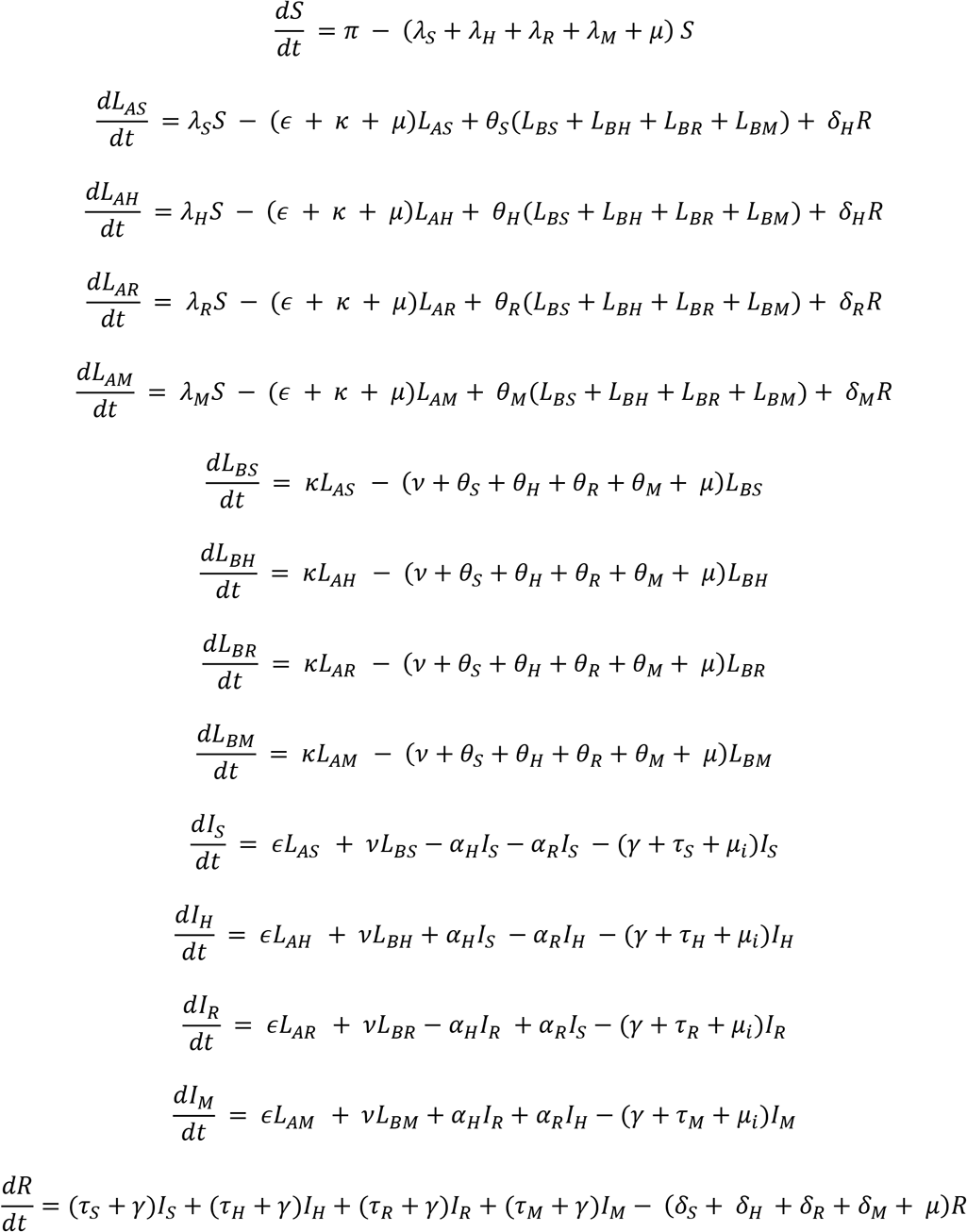

### 2.2 Parameter Estimation

An adaptive Metropolis algorithm was used to estimate model parameters, including drug resistance amplification rates. Parameters can be categorised as universal, country-specific and time-variant parameters, as presented in Table 1.

**Table 1:** Epidemiological parameters used for calibrating the model and their prior distribution ranges.

A) Universal parameters
| Parameter | Value | Prior distribution | Source |
| --- | --- | --- | --- |
| Early progression ($\epsilon$) (year<sup>-1</sup>) | 0.401775 | Uniform [ 0.1 – 0.8] | [16] |
| Transition to late latency ($\kappa$) (year<sup>-1</sup>) | 3.6525 | Uniform [1.0 – 7.0] | [16] |
| Reactivation ($\nu$) (year<sup>-1</sup>) | 0.002008875 | Uniform [0.0009, 0.006] | [16] |
| Spontaneous recovery ($\gamma$) | 0.2 | Gamma [0.16, 0.29], mode = 0.20 | [17] |
| Natural mortality ($\mu$) (year<sup>-1</sup>) | 0.0142 | | |
| TB-specific mortality ($\mu_i$) (year<sup>-1</sup>) | 0.2 | Gamma [0.06, 1.06], mode = 0.08 | [17] |
| Relative risk of reinfection rate once infected (year<sup>-1</sup>) | 0.21 | - | [18] |

B) Country-specific and time-variant parameters (used for model calibration)
| Parameter | Value | Prior distribution | Source |
| --- | --- | --- | --- |
| Transmission rate, Philippines ($\beta_{\text{Philippines}}$) | 10 | Uniform [20 - 35] | Assumption |
| Transmission rate, Viet Nam ($\beta_{\text{VietNam}}$) | 10 | Uniform [10 - 22] | Assumption |
| TSR for DS-TB (Philippines) | 0.85 | - | WHO |
| Relative TSR MDR-TB (Philippines) | 0.75 | - | Assumption |
| Relative TSR INH-R TB (Philippines) | 0.65 | - | Assumption |
| Relative TSR RIF-R TB (Philippines) | 0.70 | - | Assumption |
| TSR for DS-TB (Viet Nam) | 0.90 | - | WHO |
| Relative TSR MDR-TB (Viet Nam) | 0.60 | - | Assumption |
| Relative TSR INH-R TB (Viet Nam) | 0.65 | - | Assumption |
| Relative TSR RIF-R TB (Viet Nam) | 0.75 | - | Assumption |
| Fitness cost of INH-R TB strain | 0.90 | Uniform [0.50 – 1.20] | [19], [20] |
| Fitness cost of RIF-R TB strain | 0.70 | Uniform [0.50 – 1.20] | [21], [12] |
| Fitness cost of MDR-TB strain | 0.50 | Uniform [0.50 – 0.99] | [22] |
| Proportion of failures developing RIF-R TB ($\rho_R$) | 0.6 | Uniform [0.01 – 0.99] | Assumption |
| Proportion of failures developing INH-R TB ($\rho_H$) | 0.01 | Uniform [0.01 – 0.99] | Assumption |
| Relative risk of reinfection once recovered | 0.50 | Uniform [0.50 – 1.50] | Assumption |
| CDR start time | 1950 | Uniform [1950 -1970] | Assumption |
| CDR final value | 0.50 | Uniform [0.30 – 0.80] | Assumption |
| TSR start time | 1960 | Uniform [1960 -1980] | Assumption |

(\*TSR – Treatment success rate and CDR – Case detection rate)

#### Universal parameters

From the literature we gathered information on disease-specific and epidemiological parameters to optimise and calibrate the *Mtb* transmission model. We considered these parameters to be universal to all TB settings and so assigned the same values for all strains and settings (Table 1A).

#### Defining time-variant model processes

To capture the rise of drug resistance over time accurately, we allowed the case detection rate (CDR, a proportion) and treatment rate (τ) to vary over time. People diagnosed with active TB are commenced on treatment upon identification and move from the infectious compartments (I, I_H_, I_R_ and I_M_) to the recovered compartment (R) as they are cured. The transition from the infectious to the recovered compartment is represented using the parameter “τ”. τ is dependent on the TB detection rate “d” and the treatment success rate (TSR) and is mathematically expressed as -

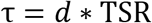

where, “d” is the TB detection rate and is defined as,

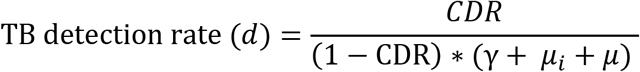

TSR is the probability of a person being tested initially and eventually put on treatment to be cured or simply put the probability of treatment success. This parameter was varied by strain but fixed over time. It is irrelevant to have a TSR value before the case detection process began; hence the TSR was fixed to a standard value. We used a sigmoidal function to model the time-variant CDR with a fixed starting time (1950), but a variable final/maximum value (e.g. 50%, as illustrated in Figure 2). By defining a scalable final CDR, we allowed flexibility in simulating the historical dynamics of TB control in the countries considered.

**Figure 2:**
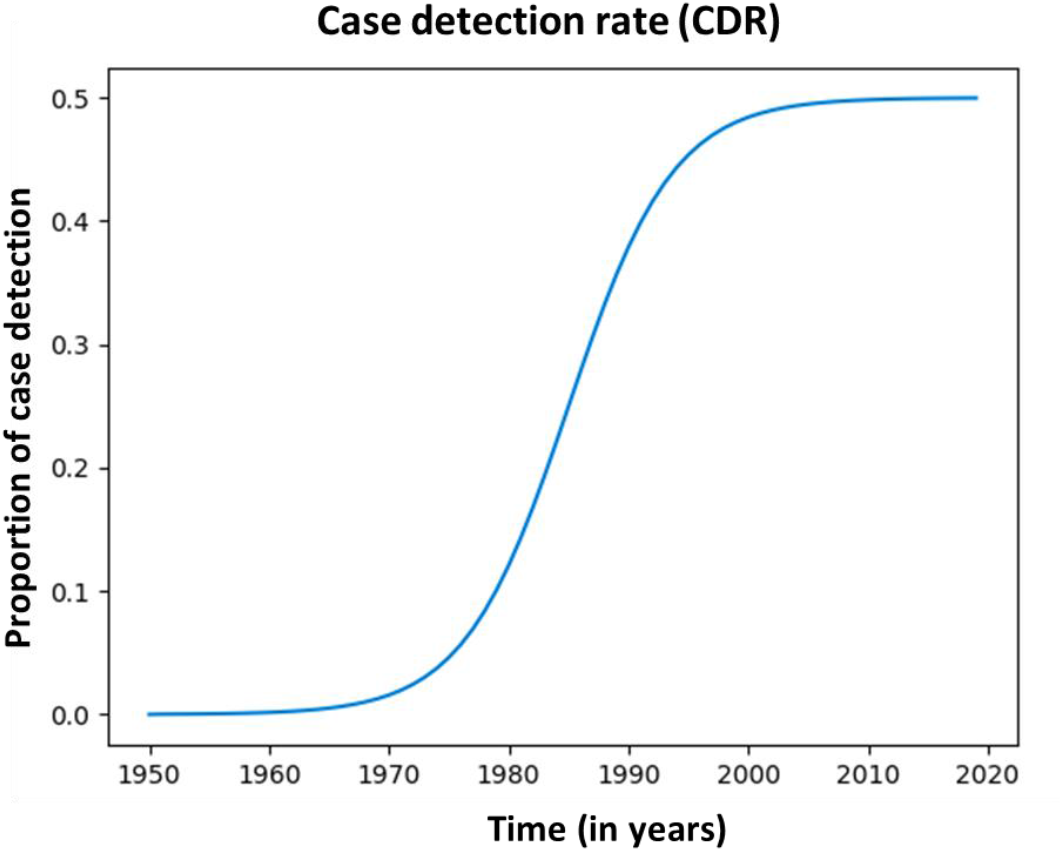
Time-variant case detection rate (CDR) (starting point=1950 and final value=50%).

#### Defining the amplification rate

Treatment for tuberculosis begins once TB is correctly diagnosed. Treatment then proceeds and may result in three possible outcomes: death, successful treatment or treatment failure. Treatment failure can further be associated with new acquisition of resistance to one additional drug that was not previously present in the infecting organism. INH and RIF are part of the standard regimen for the treatment of drug-susceptible strains. Gain in resistance to either INH or RIF is represented using amplification rates αH or αR respectively in the model. Mathematical representation of INH and RIF mono resistant amplification is shown as -

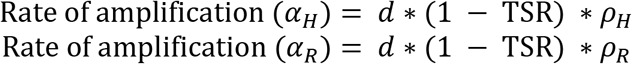

where,

*ρ_H_* = Proportion of previously INH-susceptible individuals that acquire resistance on treatment failure, and

*ρ_R_* = Proportion of previously RIF-susceptible individuals that acquire resistance on treatment failure

### 2.3 Model calibration to prevalence and notification data

#### Prevalence data

The model presented above was calibrated to individual country data. To measure the burden of TB disease, Viet Nam conducted its first nationwide population-based TB prevalence survey from 2006-2007 [23] and its second national TB prevalence survey from 2017-2018 [24]. To-date, the Philippines has conducted four national TB prevalence surveys. We calibrated our model to the third and fourth TB nationwide prevalence surveys conducted in 2007 [25] and 2016 respectively [26]. The weighted prevalence of bacteriologically confirmed TB is summarized below in Table 2. Viet Nam conducted its fourth National Anti-tuberculosis Drug Resistance Survey in 2011; reports for the first nationwide TB drug resistance survey for Philippines was published in 2009 [27] and the second in 2016 [26].

**Table 2:** Summaries of prevalence survey results and drug resistance survey data for Philippines and Viet Nam.

**A) TB Prevalence data**
| Country | Year | TB prevalence (per 100, 000) | 95% CI | Source |
| --- | --- | --- | --- | --- |
| Viet Nam | 2006-2007 | 307.2 | 248.8 – 365.6 | [23] |
|  | 2017-2018 | 322 | 260 – 399 | [24] |
| Philippines | 2007 | 660 | 510-810 | [25] |
|  | 2016 | 1159 | 1016-1301 | [26] |

**B) Drug resistance data**
| Country | Drug resistance | Year | Drug resistance (%) | 95% CI | Source |
| --- | --- | --- | --- | --- | --- |
| Viet Nam | Isoniazid mono resistance | 2011 | 14.86 | 12.15 – 17.56 | [28] |
|  | Rifampicin mono resistance | 2011 | 0.23 | 0.1 – 0.35 | [28] |
|  | MDR-TB | 2011 | 6.93 | 4.22 – 9.63 | [28] |
| Philippines | Isoniazid mono resistance | 2009 | 9.44 | 7.95 - 10.92 | [27] |
|  | Rifampicin mono resistance | 2009 | 1.008 | 0.71 - 1.304 | [27] |
|  | MDR-TB | 2009 | 5.8 | 4.3 – 7.5 | [27] |
| Philippines | Isoniazid mono resistance | 2016 | 12.43 | 11.1 - 13.75 | [26] |
|  | Rifampicin mono resistance | 2016 | 0.82 | 0.44 - 1.19 | [26] |
|  | MDR-TB | 2016 | 3.35 | 2.53 - 4.41 | [26] |

#### Notification data

We used WHO-reported TB notifications as a calibration target for both models. For Viet Nam, in 2018, 102,171 cases were notified and for Philippines 382,543 cases were notified and we calibrated to the per capita notification rates corresponding to these values.

#### Uncertainty analysis

Once we defined the parameters in our model, we next reviewed literature for information on the prior distributions of uncertain parameters (Table 1B).

The adaptive Metropolis algorithm [29] was used to generate samples from the posterior distribution of the parameters from 25,000 iterations for Philippines and Viet Nam. The primary estimates are reported as the posterior median value for all parameters of interest such as amplification rates (α_H_ and α_R_), CDR, fitness cost of each modelled strain and the relative risk of infection once recovered (δ). The intervals reported are obtained by calculating the 25^th^ and 75^th^ percentile of each parameter’s posterior distribution. Programming was done in Python 3.7.3 and all code and associated data are publicly available on GitHub (github.com/malanchak/AuTuMN).

## Results

We applied a Bayesian framework to infer the posterior distributions for the parameters in the model. Uniform priors were adopted, and tools of Bayesian inference applied.

### Calibration and optimization of the model

Figure 3A, 3B, 3C and 3D shows the model being optimized and calibrated to prevalence of infection data and reported INH-R, RIF-R and MDR levels for the high DR-TB settings, Philippines and Viet Nam respectively.

**Figure 3:**
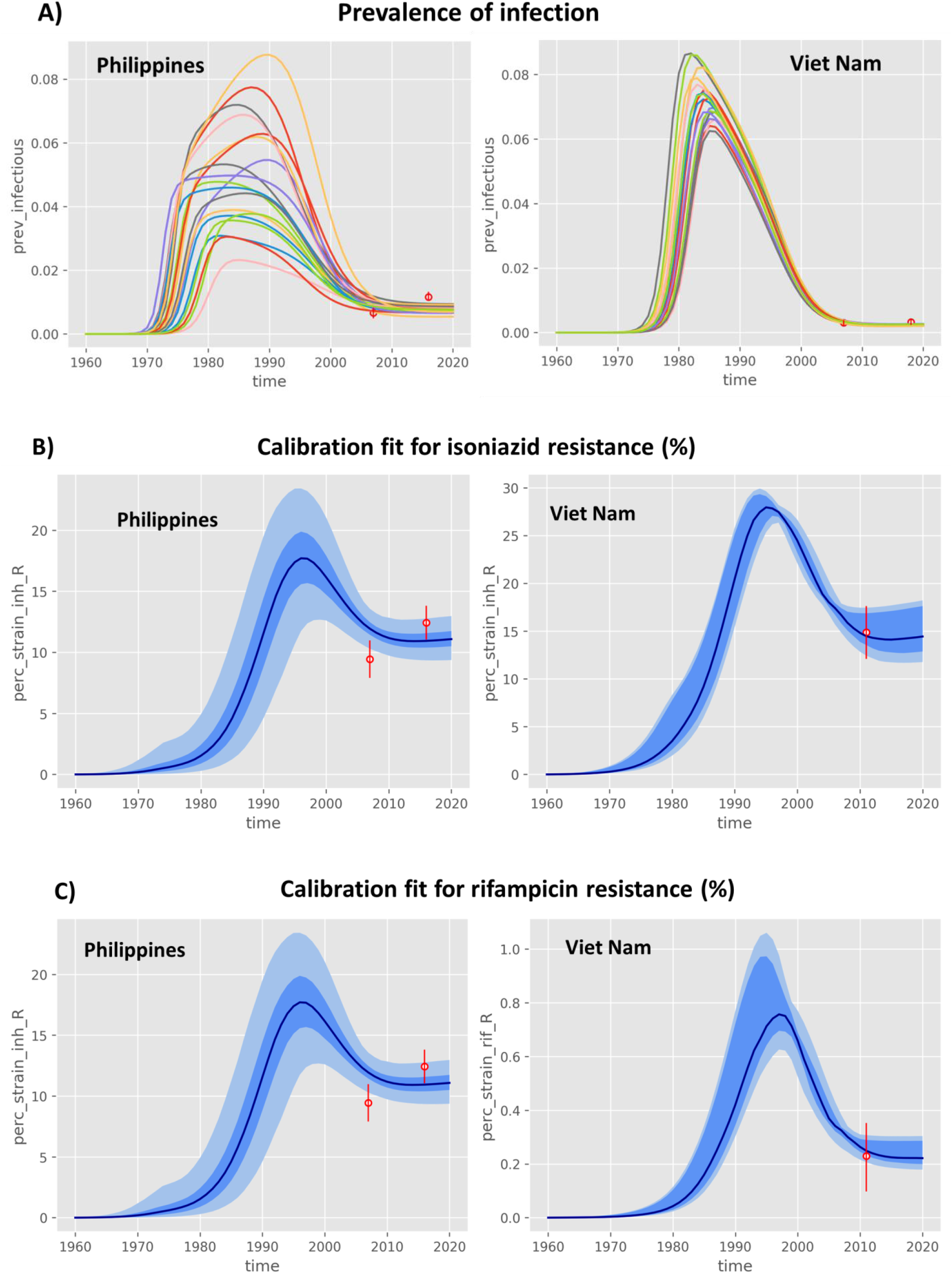
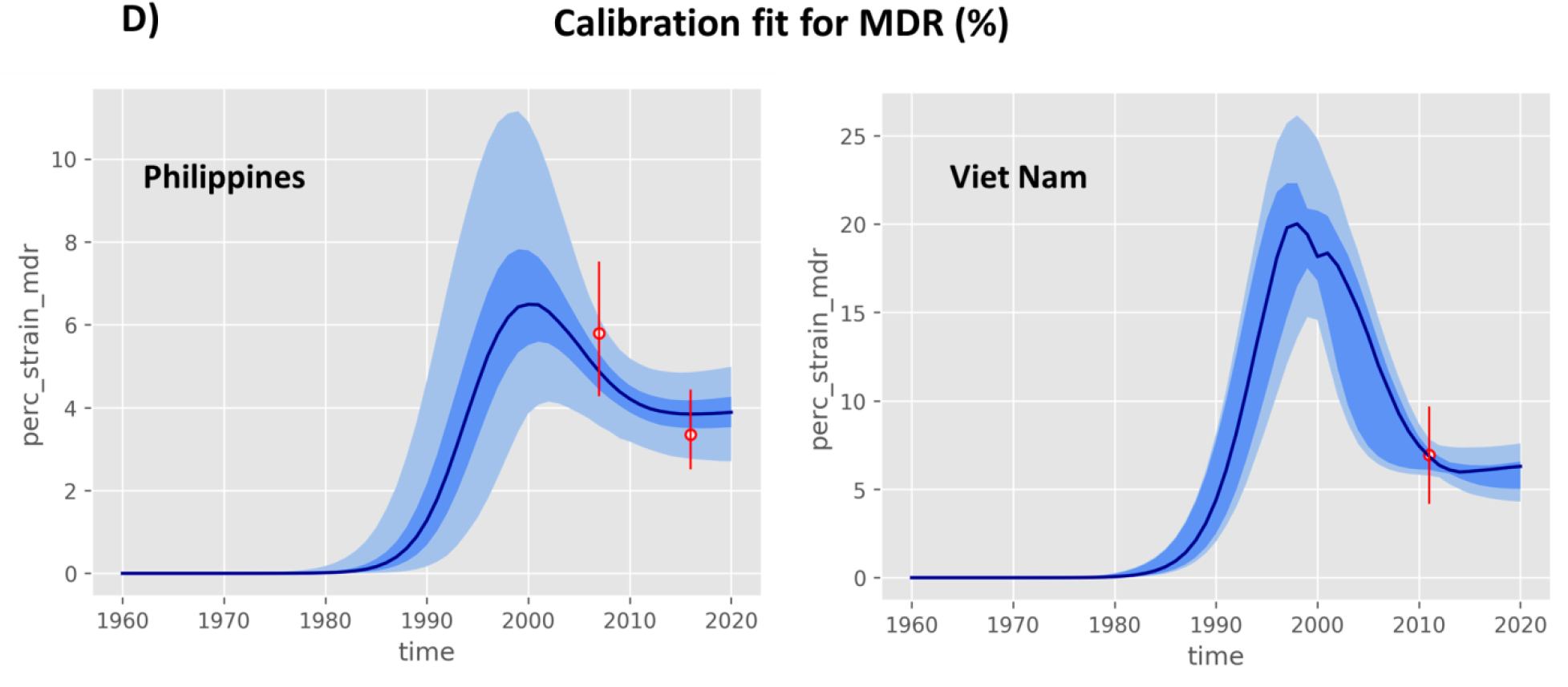
Model calibration. A) Prevalence of infection B) Isoniazid mono resistance C) Rifampicin mono resistance and D) MDR-TB

Table 3 shows the posterior distributions of all parameters.

**Table 3:** Posterior distribution of parameters obtained using the Bayesian analysis.

| Parameter | Median | 50% CI (Credible interval) | Std. Dev. | Country |
| --- | --- | --- | --- | --- |
| Proportion of previously INH-susceptible individuals that acquire resistance on treatment failure | 0.75 | 0.64 – 0.85 | 0.13 | Philippines |
|  | 0.55 | 0.39 – 0.63 | 0.01 | Viet Nam |
| Proportion of previously RIF-susceptible individuals that acquire resistance on treatment failure | 0.05 | 0.04 – 0.06 | 0.02 | Philippines |
|  | 0.011 | 0.010 – 0.012 | 0.01 | Viet Nam |
| CDR final/maximum value | 0.50 | 0.45 – 0.54 | 0.06 | Philippines |
|  | 0.62 | 0.58 – 0.73 | 0.07 | Viet Nam |
| Relative fitness of isoniazid-resistant TB | 1.04 | 0.94 – 1.11 | 0.10 | Philippines |
|  | 1.00 | 0.97 – 1.07 | 0.05 | Viet Nam |
| Relative fitness of rifampicin-resistant TB | 0.83 | 0.75 – 0.91 | 0.12 | Philippines |
|  | 0.71 | 0.68 – 0.77 | 0.04 | Viet Nam |
| Relative fitness of MDR-TB | 0.78 | 0.68 – 0.87 | 0.12 | Philippines |
|  | 0.57 | 0.54 – 0.67 | 0.06 | Viet Nam |
| Effective contact rate ($\beta$) (year<sup>-1</sup>) | 27 | 25 - 29 | 3.02 | Philippines |
|  | 11 | 10 - 13 | 1.22 | Viet Nam |
| Relative risk of re-infection after recovery ($\delta$) (year<sup>-1</sup>) | 0.82 | 0.68 – 0.95 | 0.19 | Philippines |
|  | 0.69 | 0.53 – 0.90 | 0.18 | Viet Nam |
| Rate of rapid progression ($\epsilon$) (year<sup>-1</sup>) | 0.38 | 0.31 – 0.48 | 0.12 | Philippines |
|  | 0.37 | 0.35 – 0.39 | 0.03 | Viet Nam |
| Rate of transition towards late latency ($\kappa$) (year<sup>-1</sup>) | 5.94 | 5.18 – 6.51 | 0.91 | Philippines |
|  | 2.63 | 2.11 – 2.78 | 0.37 | Viet Nam |
| Rate of re-activation ($\nu$) (year<sup>-1</sup>) | 0.0018 | 0.0016 – 0.0021 | 0.0004 | Philippines |
|  | 0.0010 | 0.0008 – 0.0013 | 0.0002 | Viet Nam |
| Self-recovery ($\gamma$) (year<sup>-1</sup>) | 0.13 | 0.10 -0.16 | 0.04 | Philippines |
|  | 0.17 | 0.13 – 0.23 | 0.065 | Viet Nam |
| Transmission rate of INH-R (%) | 56 | 42 – 68 | - | Philippines |
|  | 79 | 70 – 86 | - | Viet Nam |
| Transmission rate of RIF-R (%) | 48 | 34 – 62 | - | Philippines |
|  | 68 | 58 – 75 | - | Viet Nam |
| Transmission rate of MDR (%) | 42 | 34 – 50 | - | Philippines |
|  | 50 | 45 – 53 | - | Viet Nam |

### Drug resistance amplification and transmission

We observed higher rates of drug resistance amplification for isoniazid compared to rifampicin for both the high DR-TB incidence settings (Figure 4). The estimated proportion of isoniazid mono-resistance amplification due to treatment failure was 0.75 (0.64 – 0.85) for Philippines and 0.55 (0.39 – 0.63) for Viet Nam. The rates of rifampicin mono-resistance amplification were 0.05 (0.04 – 0.06) for Philippines and 0.011 (0.010 – 0.012) for Viet Nam. This meant approximately 75% and 55% of the people who failed treatment in Philippines and Viet Nam respectively would end up with resistance to isoniazid. Several studies performed in high DR-TB burden countries report resistance to isoniazid (INH) were very common, often in the absence of resistance to rifampicin (RIF) [30].

**Figure 4:**
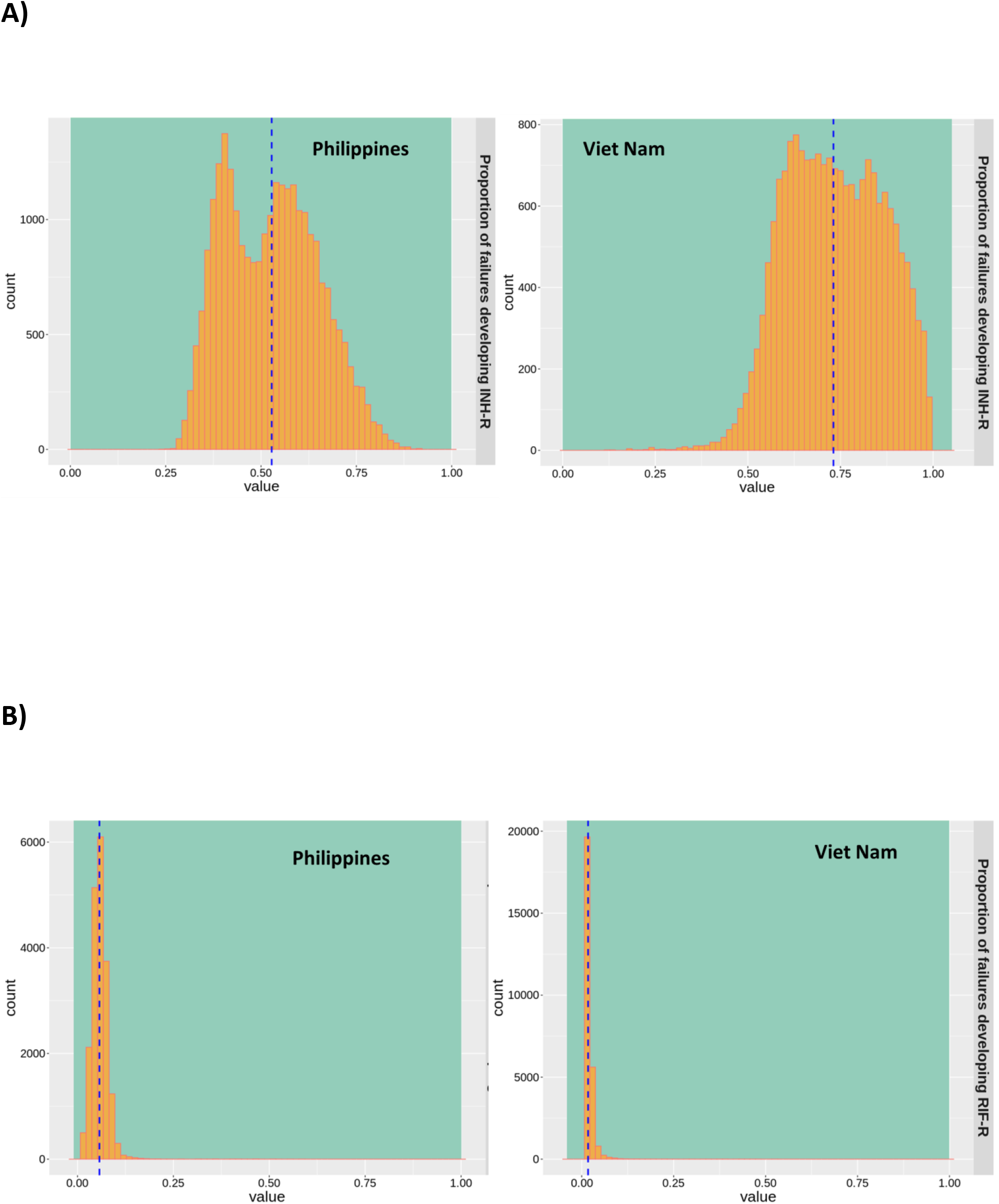
The estimated proportion of mono-drug resistance amplification for INH and RIF due to treatment failure. The histogram represents the posterior distribution of the estimates of amplification and the green background represents the prior ranges. The dashed blue line is the median of the estimates. A) Proportion of previously INH-susceptible strains that acquire resistance on treatment failure and B) Proportion of previously RIF-susceptible strains that acquire resistance on treatment failure.

With respect to estimated transmission rates, for Philippines, INH-R transmission was 56% (42 – 68), RIF-R was 48% (34 – 62) and MDR was 42% (34 – 50). In case of Viet Nam, estimated rates of INH-R transmission was 79% (70 – 86), RIF-R transmission was 68% (58 – 75) and MDR was 50% (45 – 53).

In Philippines, the incidence data for amplification from DS to INH-R (Figure S1) shows 23 per 100,000 (17 – 30) people failing treatment initially, followed by 9 per 100,000 (6 – 12) people then gain resistance to RIF and moving to the MDR compartment. Comparing this to acquiring RIF resistance first (Figure S2), we see 1.7 per 100,000 (1.14 – 2.5) people failing treatment and moving from DS to RIF-R, followed by only 0.04 (0.02 – 0.08) people gaining resistance to INH to move to the MDR compartment. A similar observation was seen for Viet Nam, the incidence data for amplification from DS to INH-R (Figure S1) shows 5 per 100,000 (3 – 8) people failing treatment followed by people 4 per 100,000 (3 – 6) gaining resistance to RIF and moving to the MDR compartment. In case of DS to RIF-R transition (Figure S2) we see 0.1 per 100,000 (0.08 – 0.2) people failing treatment initially, followed by 0.001 per 100,000 (0.0006 – 0.002) people gaining resistance to INH to move to the MDR compartment.

### Estimates for fitness cost, CDR and risk of re-infection upon recovery

The posterior estimates of fitness cost associated with INH-R strains for Philippines was 1.04 (0.94 – 1.11) and 1.00 (0.97 – 1.07) for Viet Nam. The fitness cost associated with RIF-resistance strains for Philippines was 0.83 (0.75 – 0.91) and 0.71 (0.68 – 0.77) for Viet Nam. The fitness cost associated with MDR strains in Philippines was 0.78 (0.68 – 0.87) compared to 0.57 (0.54 – 0.67) for Viet Nam.

The uncertainty analysis provided information on the relative risk of re-infection upon recovery. The estimates for Philippines was 0.82 (0.68 – 0.95) which was higher than Viet Nam, 0.69 (0.53 – 0.94). Our study also provided information on estimates of CDR with high precision for both the settings. The estimates obtained for Philippines was 0.50 (0.45 – 0.54) and for Viet Nam was 0.62 (0.58 – 0.73).

## Discussion

Accurate diagnosis of DR-TB increases the opportunity for effective treatment, thus improving the rate of cure and survival and reducing the potential for transmission. We found higher rate of transmission and resistance amplification for INH-R compared to RIF-R for both the high DR-TB settings investigated in this model. This appears to relate to current methods of DR-TB detection which emphasize RIF’s resistance identification, and so might contribute to INH mono-resistance amplification. Further research into the association between specific INH resistance mutations and differential risk of transmission will be helpful in better defining the public health impact of this effect [31].

In addition to programmatic insights, our model provides novel information on parametrizing CDR. This is important, as this parameter cannot be measured directly yet plays a significant role in informing robust mathematical model of TB transmission. Inclusion of notification data and prevalence of infection for the MCMC analysis helped in constraining the time-variant CDR parameter. The uncertainty analysis also provided information on the relative risk of re-infection upon recovery. The estimate for Philippines was higher than Viet Nam, which implies Viet Nam has been more successful in detecting re-infection cases compared to Philippines. However, estimates for both settings are high and hence, the model suggests that people in high drug resistance TB incident setting have a higher chance of re-infection following treatment [32, 33].

As with any mathematical representation our model has certain limitation. Our model is primarily built for pulmonary tuberculosis (PTB) and does not include extra-pulmonary TB data, as our primary focus was on transmission. For the same reason, this model has been parametised from adult TB data given the limited TB transmission from young children to others. We have adopted a simplified model structure to estimate rates of mono-resistance amplifications for INH and RIF and it does not capture the heterogeneity associated with TB epidemics, which may be important in some settings.

Historically, diagnosis of MDR TB has been reliant on culture-based phenotypic testing, which in high burden settings may be applied selectively, such as after treatment failure. As part of the global policy to control DR-TB, many high burden settings have pledged to deploy the molecular diagnostic assay Xpert MTB/RIF (detects resistance only in RIF), which is a nucleic acid amplification test that can be directly applied to sputum samples [34] [35]. As the presence of RIF resistance is highly predictive of MDR-TB, these policies have led to significant improvements in initiating second-line therapy [36]. However, as our work highlights, these algorithms may also be associated with selecting for and further amplifying INH resistance. Alternative molecular tools, such as the line probe assay MTBDR*plus* [37] or Xpert MTB/RIF Ultra [38], can identify both RIF and INH resistance, and may offer the programmatic advantages of rapid MDR-TB diagnosis while avoiding this secondary effect [39].

While rapid molecular diagnostics will continue to be important for programmatic adoption, it is also important to recognize that the principle of unrecognised resistance amplification demonstrated here can be repeated for any resistance not routinely addressed in diagnostic algorithms. It is therefore essential to incorporate genome sequencing into surveillance programs, to maximise the clinical and public health benefits [40]. With recent developments in next generation sequencing techniques, we have now have high-throughput diagnostic tools for the detection of DR-TB which are both fast and efficient [41]. While such tools are currently in routine use only in high resource settings, the benefits associated with these tools should be prioritized for high burden contexts to support optimal individual and program outcomes [42, 43].

## Notes

### Competing Interest Statement

The authors have declared no competing interest.

## References

1. WHO, Global Tuberculosis Report 2020. https://www.who.int/teams/global-tuberculosis-programme/tb-reports/global-tuberculosis-report-2020, 2020.

2. WHO, WHO consolidated guidelines on drug-resistant tuberculosis treatment. 2019.

3. Ragonnet, R., et al., High rates of multidrug-resistant and rifampicin-resistant tuberculosis among re-treatment cases: where do they come from? BMC Infectious Diseases, 2017. 17(1): p. 36.

4. Trauer, J.M., et al., Modular programming for tuberculosis control, the “AuTuMN” platform. BMC Infectious Diseases, 2017. 17(1): p. 546.

5. Zwerling, A., S. Shrestha, and D.W. Dowdy, Mathematical Modelling and Tuberculosis: Advances in Diagnostics and Novel Therapies. Advances in Medicine, 2015. 2015: p. 907267.

6. Fors, J., et al., Mathematical model and tool to explore shorter multi-drug therapy options for active pulmonary tuberculosis. PLoS Comput Biol, 2020. 16(8): p. e1008107.

7. Pontali, E., M.C. Raviglione, and G.B. Migliori, Regimens to treat multidrug-resistant tuberculosis: past, present and future perspectives. European Respiratory Review, 2019. 28(152): p. 190035.

8. Herrera, M., et al., Modeling the Spread of Tuberculosis in Semiclosed Communities. Computational and Mathematical Methods in Medicine, 2013. 2013: p. 648291.

9. Wikell, A., et al., Diagnostic pathways and delay among tuberculosis patients in Stockholm, Sweden: a retrospective observational study. BMC Public Health, 2019. 19(1): p. 151.

10. Naidoo, P., et al., Pathways to multidrug-resistant tuberculosis diagnosis and treatment initiation: a qualitative comparison of patients’ experiences in the era of rapid molecular diagnostic tests. BMC health services research, 2015. 15: p. 488–488.

11. Becerra, M.C., et al., Transmissibility and potential for disease progression of drug resistant *Mycobacterium tuberculosis*: prospective cohort study. BMJ, 2019. 367: p. l5894.

12. Knight, G.M., et al., The Distribution of Fitness Costs of Resistance-Conferring Mutations Is a Key Determinant for the Future Burden of Drug-Resistant Tuberculosis: A Model-Based Analysis. Clin Infect Dis, 2015. 61Suppl 3(Suppl 3): p. S147–54.

13. Karmakar, M., et al., Analysis of a Novel pncA Mutation for Susceptibility to Pyrazinamide Therapy. Am J Respir Crit Care Med, 2018. 198(4): p. 541–544.

14. Karmakar, M., et al., Structure guided prediction of Pyrazinamide resistance mutations in pncA. Sci Rep, 2020. 10(1): p. 1875.

15. Karmakar, M., et al., Empirical ways to identify novel Bedaquiline resistance mutations in AtpE. PLoS One, 2019. 14(5): p. e0217169.

16. Ragonnet, R., et al., Optimally capturing latency dynamics in models of tuberculosis transmission. Epidemics, 2017. 21: p. 39–47.

17. Ragonnet, R., et al., Revisiting the Natural History of Pulmonary Tuberculosis: a Bayesian Estimation of Natural Recovery and Mortality rates. Clin Infect Dis, 2020.

18. Trauer, J.M., et al., Risk of Active Tuberculosis in the Five Years Following Infection…15%? Chest, 2016. 149(2): p. 516–525.

19. Cohen, T., B. Sommers, and M. Murray, The effect of drug resistance on the fitness of Mycobacterium tuberculosis. Lancet Infect Dis, 2003. 3(1): p. 13–21.

20. Borrell, S. and S. Gagneux, Infectiousness, reproductive fitness and evolution of drug-resistant Mycobacterium tuberculosis. Int J Tuberc Lung Dis, 2009. 13(12): p. 1456–66.

21. Gagneux, S., Fitness cost of drug resistance in Mycobacterium tuberculosis. Clin Microbiol Infect, 2009. 15 Suppl 1: p. 66–8.

22. Cohen, T. and M. Murray, Modeling epidemics of multidrug-resistant M. tuberculosis of heterogeneous fitness. Nat Med, 2004. 10(10): p. 1117–21.

23. Hoa, N.B., et al., National survey of tuberculosis prevalence in Viet Nam. Bull World Health Organ, 2010. 88(4): p. 273–80.

24. Nguyen, H.V., et al., The second national tuberculosis prevalence survey in Vietnam. PLoS One, 2020. 15(4): p. e0232142.

25. Tupasi, T.E., et al., Significant decline in the tuberculosis burden in the Philippines ten years after initiating DOTS. Int J Tuberc Lung Dis, 2009. 13(10): p. 1224–30.

26. Department of Health, R.o.T.P., National Tuberculosis Prevalence Survey 2016 Philippines. 2016. http://www.ntp.doh.gov.ph/downloads/publications/Philippines_2016%20National%20TB%20Prevalence%20Survey_March2018.pdf.

27. Nationwide drug resistance survey of tuberculosis in the Philippines. Int J Tuberc Lung Dis, 2009. 13(4): p. 500–7.

28. Nhung, N.V., et al., The fourth national anti-tuberculosis drug resistance survey in Viet Nam. Int J Tuberc Lung Dis, 2015. 19(6): p. 670–5.

29. Haario, H., E. Saksman, and J. Tamminen, An Adaptive Metropolis Algorithm. Bernoulli, 2001. 7(2): p. 223–242.

30. Kumar, P., et al., High degree of multi-drug resistance and hetero-resistance in pulmonary TB patients from Punjab state of India. Tuberculosis (Edinb), 2014. 94(1): p. 73–80.

31. Fregonese, F., et al., Comparison of different treatments for isoniazid-resistant tuberculosis: an individual patient data meta-analysis. Lancet Respir Med, 2018. 6(4): p. 265–275.

32. Yew, W.W. and C.C. Leung, Are some people not safer after successful treatment of tuberculosis? Am J Respir Crit Care Med, 2005. 171(12): p. 1324–5.

33. Sonnenberg, P., et al., HIV-1 and recurrence, relapse, and reinfection of tuberculosis after cure: a cohort study in South African mineworkers. Lancet, 2001. 358(9294): p. 1687–93.

34. Tsara, V., E. Serasli, and P. Christaki, Problems in diagnosis and treatment of tuberculosis infection. Hippokratia, 2009. 13(1): p. 20–2.

35. Campbell, E.A., et al., Structural mechanism for rifampicin inhibition of bacterial rna polymerase. Cell, 2001. 104(6): p. 901–12.

36. Dlamini, M.T., et al., Whole genome sequencing for drug-resistant tuberculosis management in South Africa: What gaps would this address and what are the challenges to implementation? Journal of clinical tuberculosis and other mycobacterial diseases, 2019. 16: p. 100115–100115.

37. Nathavitharana, R.R., et al., Multicenter Noninferiority Evaluation of Hain GenoType MTBDRplus Version 2 and Nipro NTM+MDRTB Line Probe Assays for Detection of Rifampin and Isoniazid Resistance. J Clin Microbiol, 2016. 54(6): p. 1624–1630.

38. Chakravorty, S., et al., Detection of Isoniazid-, Fluoroquinolone-, Amikacin-, and Kanamycin-Resistant Tuberculosis in an Automated, Multiplexed 10-Color Assay Suitable for Point-of-Care Use. J Clin Microbiol, 2017. 55(1): p. 183–198.

39. Talbot, E.A. and M. Pai, Tackling drug-resistant tuberculosis: we need a critical synergy of product and process innovations. Int J Tuberc Lung Dis, 2019. 23(7): p. 774–782.

40. Dunstan, S.J., D.A. Williamson, and J.T. Denholm, Understanding the global tuberculosis epidemic: moving towards routine whole-genome sequencing. Int J Tuberc Lung Dis, 2019. 23(12): p. 1241–1242.

41. Mahomed, S., et al., Whole genome sequencing for the management of drug-resistant TB in low income high TB burden settings: Challenges and implications. Tuberculosis (Edinb), 2017. 107: p. 137–143.

42. Luo, T., et al., Whole-genome sequencing to detect recent transmission of Mycobacterium tuberculosis in settings with a high burden of tuberculosis. Tuberculosis (Edinb), 2014. 94(4): p. 434–40.

43. Sulis, G. and M. Pai, Isoniazid-resistant tuberculosis: A problem we can no longer ignore. PLoS Med, 2020. 17(1): p. e1003023.

