## Supplementary Data for "Modelling tuberculosis drug resistance amplification rates in high-burden settings"

### S1: Modelled incidence rate of de novo acquired resistant TB. The first plot shows incidence rates from DS-TB to INH-R-TB followed by INH-R-TB to MDR-TB in the second plot.

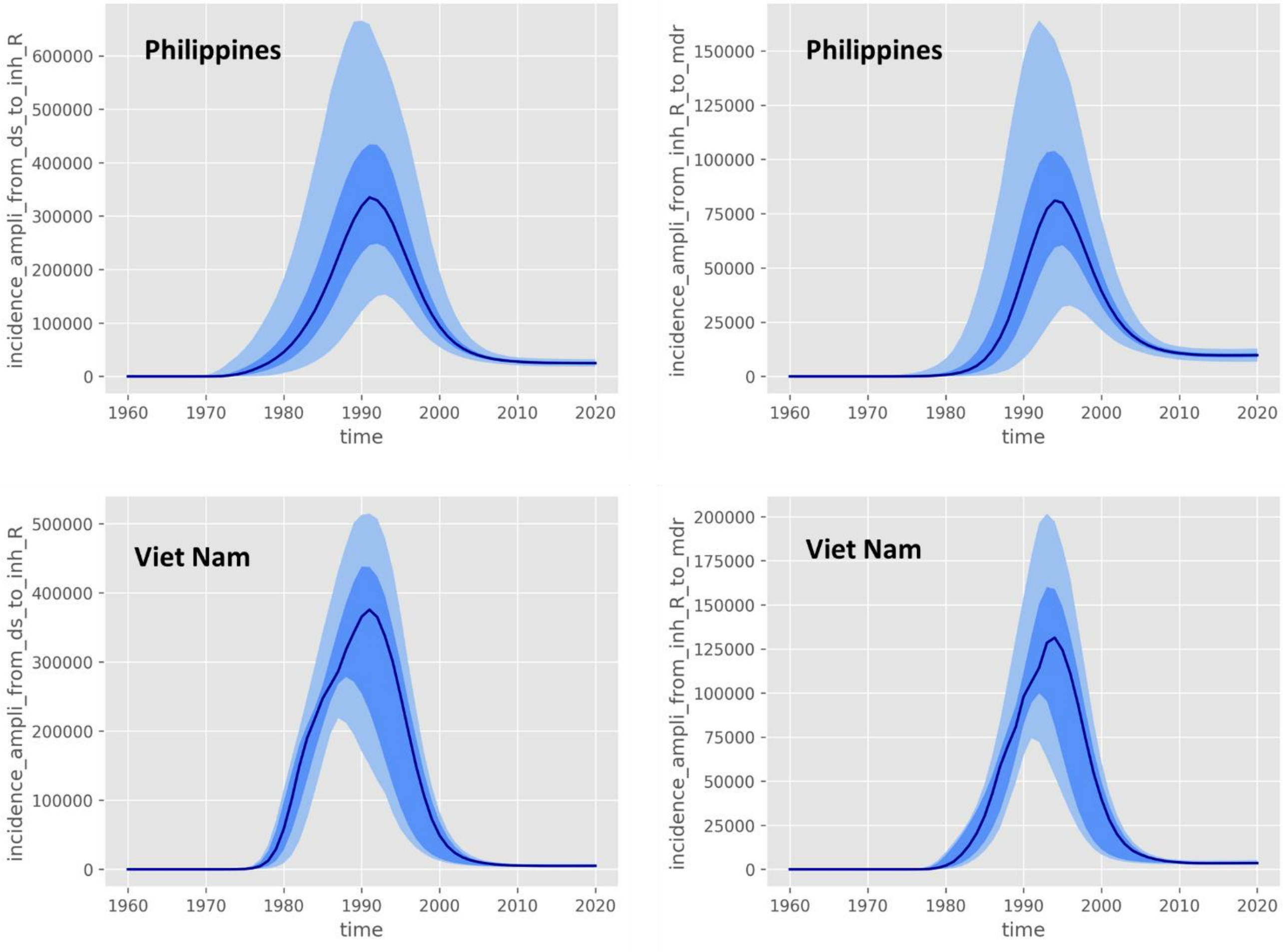

**S2: Modelled incidence rate of de novo acquired resistant TB. The first plot shows incidence rates from DS-TB to RIF-R-TB followed by RIF-R-TB to MDR-TB in the second plot.**

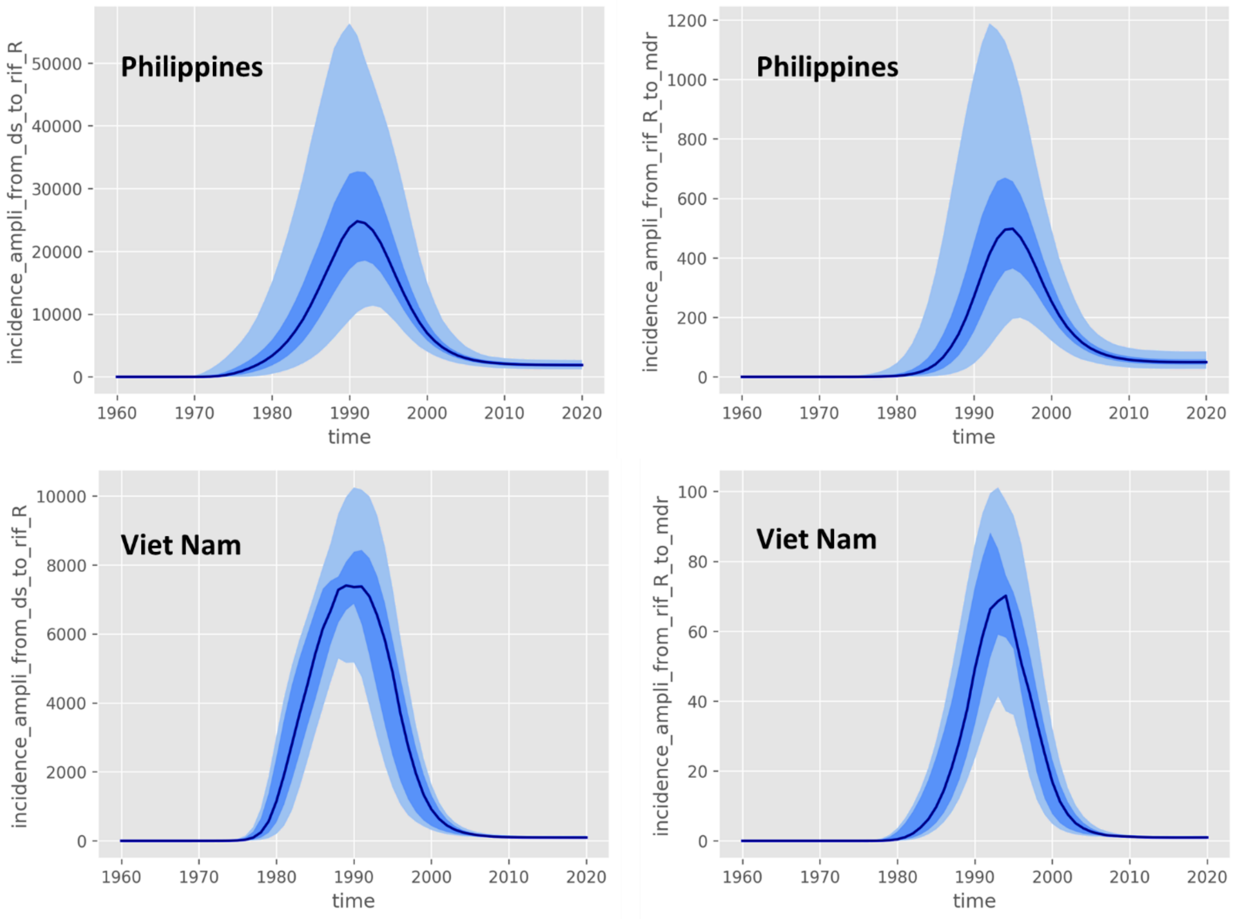
